# Associations between polygenic liability for schizophrenia and level of psychosis and mood-incongruence in bipolar disorder

**DOI:** 10.1101/160119

**Authors:** Judith Allardyce, Ganna Leonenko, Marian Hamshere, Antonio F. Pardiñas, Liz Forty, Sarah Knott, Katherine-Gordon Smith, David J. Porteus, Caroline Haywood, Arianna Di Florio, Lisa Jones, Andrew M. McIntosh, Michael J. Owen, Peter Holmans, James T.R. Walters, Nicholas Craddock, Ian Jones, Michael C. O’Donovan, Valentina Escott-Price

**Affiliations:** MRC Centre for Neuropsychiatric Genetics and Genomics, Institute of Psychological Medicine and Clinical Neurosciences, School of Medicine, Cardiff University, Cardiff, UK; Department of Psychological Medicine, University of Worcester, Worcester, UK; Medical Genetics Section, Centre for Genomic and Experimental Medicine, Institute of Genetics and Molecular Medicine, University of Edinburgh, Edinburgh, UK; Division of Psychiatry, University of Edinburgh, Edinburgh, UK

## Abstract

**Importance:** Bipolar disorder (BD) overlaps schizophrenia in its clinical presentation and genetic liability. Alternative approaches to patient stratification beyond current diagnostic categories are needed to understand the underlying disease processes/mechanisms.

**Objectives:** To investigate the relationship between common-variant liability for schizophrenia, indexed by polygenic risk scores (PRS) and psychotic presentations of BD, using clinical descriptions which consider both occurrence and level of mood-incongruent psychotic features.

**Design:** Case-control design: using multinomial logistic regression, to estimate differential associations of PRS across categories of cases and controls.

**Settings & Participants:** 4399 BDcases, mean [sd] age-at-interview 46[12] years, of which 2966 were woman (67%) from the BD Research Network (BDRN) were included in the final analyses, with data for 4976 schizophrenia cases and 9012 controls from the Type-1 diabetes genetics consortium and Generation Scotland included for comparison.

**Exposure:** Standardised PRS, calculated using alleles with an association p-value threshold < 0.05 in the second Psychiatric Genomics Consortium genome-wide association study of schizophrenia, adjusted for the first 10 population principal components and genotyping-platform.

**Main outcome measure:** Multinomial logit models estimated PRS associations with BD stratified by (1) Research Diagnostic Criteria (RDC) BD subtypes (2) Lifetime occurrence of psychosis.(3) Lifetime mood-incongruent psychotic features and (4) ordinal logistic regression examined PRS associations across levels of mood-incongruence. Ratings were derived from the Schedule for Clinical Assessment in Neuropsychiatry interview (SCAN) and the Bipolar Affective Disorder Dimension Scale (BADDS).

**Results:** Across clinical phenotypes, there was an exposure-response gradient with the strongest PRS association for schizophrenia (RR=1.94, (95% C.I. 1.86, 2.01)), then schizoaffective BD (RR=1.37, (95% C.I. 1.22, 1.54)), BD I (RR= 1.30, (95% C.I. 1.24, 1.36)) and BD II (RR=1.04, (95% C.I. 0.97, 1.11)). Within BD cases, there was an effect gradient, indexed by the nature of psychosis, with prominent mood-incongruent psychotic features having the strongest association (RR=1.46, (95% C.I. 1.36, 1.57)), followed by mood-congruent psychosis (RR= 1.24, (95% C.I. 1.17, 1.33)) and lastly, BD cases with no history of psychosis (RR= 1.09, (95% C.I. 1.04, 1.15)).

**Conclusion:** We show for the first time a polygenic-risk gradient, across schizophrenia and bipolar disorder, indexed by the occurrence and level of mood-incongruent psychotic symptoms.

## Key Points

**Question**: what is the relationship between schizophrenia related polygenic liability and the occurrence and level of mood-incongruence of psychotic symptoms in bipolar disorder (BD)?

**Findings**: in this case-control study including 4436 BD cases, 4976 schizophrenia cases and 9012 controls, there was an exposure-response gradient of polygenic risk: Schizophrenia > BD with prominent mood-incongruent psychotic features > BD with mood-congruent psychotic features > BD with no psychosis, all differential associations were statistically-significant.

**Meaning**: A gradient of genetic liability across schizophrenia and bipolar disorder, indexed by the occurrence of psychosis and level of mood-incongruence has been shown for the first time.

## Introduction

Although classified as a discrete diagnostic category^1-3^, bipolar disorder (BD) overlaps considerably with schizophrenia (SCZ) in both its clinical presentation ^4–13^ and genetic liability ^14–22^. BD is a phenomenologically heterogeneous construct and within the diagnostic category, individuals may have quite different symptom profiles. It has been proposed, that this clinical heterogeneity indicates underlying aetiological heterogeneity and the degree of clinical similarity between BD and SCZ reflects, overlapping alleles which selectively influence specific, shared clinical characteristics, rather than the global risk for the disorders ^23–25^.

Delusions and hallucinations are common in BD ^26,27^ with around one third of all psychotic features judged to be mood-incongruent^28,29^. Mood-incongruent psychotic features, are associated with poorer prognosis, poor lithium-response and are qualitatively similar to the prototypic symptoms of SCZ ^30–32^, suggesting that BD with psychosis and particularly mood-incongruent psychotic features, may specify a sub group/stratum with stronger aetiological links to SCZ. Stratified linkage and candidate-gene studies of BD associations with chromosomal regions and genes implicated in SCZ, show stronger effects in psychosis and mood-incongruent subsamples ^33–36^ providing some support for this causal heterogeneity hypothesis, however lack of consistency in earlier linkage and candidate-gene studies renders the overall support weak.

Recently, genome-wide association studies (GWAS) have found a substantial polygenic component to both BD and SCZ risk, with a large proportion of their genetic variance explained by common alleles, partially shared across the two disorders^20^. Polygenic-risk can be calculated for individuals, with a single summary measure: the polygenic risk score (PRS), which allows us to examine the genetic basis of symptom domains, within and across the two disorders ^37–39^ with greater power than the historical linkage and candidate-gene approaches. PRS-SCZ differentiate BD from controls^20,40^ and there are differential associations across subtypes with schizoaffective bipolar disorder (SABD) (intermediate subtype, characterised by admixture of SCZ and BD symptoms) having a relatively larger burden of SCZ risk, compared to other BD subtypes^15,41^. To date, lack of power in well phenotyped samples has hindered fine-scale examination of the relationship between SCZ polygenic-risk and psychotic symptoms in BD.

We aimed to examine the relationship between polygenic liability for SCZ and psychotic presentations of BD using PRS generated from the most powerful SCZ-GWAS discovery set available, currently^21^. Measures relevant to the occurrence and nature of psychotic symptoms were considered. We hypothesised BD with psychosis would be associated with higher polygenic-risk for SCZ and this association would be stronger when mood-incongruent psychotic features were present, given their phenotypic similarity to the psychotic symptoms of prototypic SCZ.

## Methods

### Sample Ascertainment

#### Bipolar Disorder sample

4436 cases of BD with deep phenotypic information, European ancestry, domicile in the UK, collected between 2000 - 2013 were available via the UK BD Research Network (BDRN) using recruitment methods reported previously ^15,42^,^43^. The sample has 1399 cases not included in prior BDRN publications ^15,41^. All participants were assessed using a consistent protocol which included the Schedule for Clinical Assessment in Neuropsychiatry interview (SCAN) ^44^ administered by trained research psychologists and psychiatrists, with very good to excellent inter-rater reliability for all domains of psychopathology ^45^. Using information from the SCAN and casenote review, the Operational Criteria Checklist (OPCRIT) ^46^ was completed. Research Diagnostic Criteria (RDC) ^3^diagnoses, which differentiate individuals on the basis of the their pattern of mood and psychotic symptoms better ^41^ than either DSM^2^ or ICD-10^1^, were made using the consensus lifetime best-estimate method, informed by all available information^47^.

#### Schizophrenia sample

To allow comparison of BD with SCZ, we included a subset (N=4976) of the CLOZUK sample, collected via the Zapronex^®^ Treatment Access System as detailed in a previous report^48^, All were prescribed clozapine for treatment resistant SCZ (TRS) and are independent of, and unrelated (pi-hat <0.2) to individuals in the discovery GWAS^21^. In principle, TRS may carry higher polygenic-risk burden, however PRS in CLOZUK are similar to the other SCZ samples used by the Psychiatric Genomics Consortium^21^.

#### Control Samples

The controls came from two UK sources: the Type-1 diabetes genetics consortium (TIDGC) (n = 2,532) are unscreened controls, recruited through the 1958 birth-cohort ^49^ and the other is a subsample of the Generation Scotland (n = 6,480) study, screened for psychiatric disorders^50^. Controls are unrelated (pi-hat < 0.2) to individuals in the PGC-SCZ discovery set, and were matched ancestrally to our case datasets ^48^

All samples have appropriate ethics approvals.

### Genotyping, quality control (QC), phasing and imputation

#### Bipolar cases

Genotypic data for the BD cases were processed in 3 batches, each on a different platform. To mitigate against potential bias from batch effects^51^, stringent QC was performed on each platform separately prior to merging. Single nucleotide polymorphisms (SNPs) were excluded if the call rate was < 98%, MAF was < 0.01 or they deviated from HWE at p < l×10^6^. Individuals were excluded if they had minimal or excessive autosomal homozygosity (|F|>0.1), high pairwise relatedness (pi-hat > 0.2) or mismatch between recorded and genotypic sex. Following QC, the data for each platform were phased using SHAPEIT^52^ and imputed with IMPUTE2 ^53^, using the 1000 Genomes reference panel (Phase3,2014). Imputed data were converted into the most probable genotypes (probability >0.9) and merged on shared SNPs. 4399 BD cases remained after QC.

### CLOZUK cases and Controls

The CLOZUK and control samples had been though strict QC separately, before being phased and imputed simultaneously as part of a larger SCZ study^48^

#### Merging BD, CLOZUK and control imputed genotypic datasets

After excluding SNPs with stand ambiguity; BD, CLOZUK and control samples were merged and the imputed markers underwent a second QC Alter^51^, excluding SNPs with; missingness in >5% of individuals, (INFO) <0.8, MAF <0.01 or deviation from HWE atp < 1×10^-6^.

### Principal Component Analysis

To adjust for potential confounding from population structure, we performed PCA using PLINK v1.9, after LD pruning and frequency filtering the SNPs from the merged sample, keeping the eigenvectors for the first 10 principal components (PCs) to use as covariates in the association analysis.

### Polygenic Risk Scores (PRS)

We generated PRS^20^, using the 2014 PGC-SCZ meta-analysis as our discovery set^21^ calculated for each individual, based on a set of alleles with association p-values < 0.05. This decision was informed by the PGC leave one-cohort-out PRS analyses, for all SNP selection p-value thresholds, which found the median and mode of the cut-off = 0.05. This represents the association that best optimises the balance of false and true risk alleles, at the current discovery sample size^21^. The most informative and independent markers were selected to minimise statistical noise where possible, using p-value informed clumping, atr^2^ <0.2 with 1MB windows and by excluding the extended MHC (Chr6: position 25-35MB) because of its complex LD structure.

### Outcome measure of lifetime psychosis & mood incongruence

#### Subtypes of BD

RDC subtypes were used as categorical outcomes in case-control analyses. The RDC^3^ and Diagnostic and Statistical Manual of Mental Disorders (DSM) ^2^, though not the ICD-

10 Classification of Mental and Behavioural Disorder (ICD-10) ^1^, subdivides BD into bipolar I disorder (BD I) and bipolar II disorder (BD II) depending on the nature of the mood states; mania in (BP I) and hypomania in (BP II). All classification systems recognise SABD. Psychotic symptoms are most prominent in SABD, then BD I, and least prominent in BD II ^54,55^.

#### The Bipolar Affective Disorder Dimension Scale

Outcome measures were generated from The Bipolar Affective Disorder Scale (BADDS) Psychosis (P) and mood-incongruence (I) subscales, which provide an ordered (not necessarily linear) measure of lifetime symptom domain severity^56^. An inter-rater reliability exercise for this sample demonstrates excellent interclass correlation: (P) 0.91 and (I) 0.89.

1) A binary categorical outcome measure for lifetime occurrence of psychosis defined as an unambiguous episode of positive and/or disorganised psychotic symptoms, generated by dichotomising the (P) domain scale at a score > 9 ^56^.
2) A binary categorical outcome measure for lifetime occurrence of predominant mood-incongruent psychotic features (high v low prominence of mood-incongruence), generated by dichotomising the (I) domain scale at a score >19.
3) An ordinal measure of mood-incongruent psychotic features which assesses the overall balance between mood-congruent and mood-incongruent psychosis across the lifetime, rated using all available information according to BDRN protocol (E supplement: Note 1)

### Statistical Analysis

A multinomial logit model (MNLM) was used to estimate differential associations of standardised PRS, adjusted for the first 10 PCs and genotyping-platform, across categories of cases and controls. We report the estimated coefficients transformed to relative risk-ratios (RR), defined as the exponentiated regression coefficient PRS association across levels of mood-incongruent psychotic features using ordinal logistic regression was also estimated. To examine whether SABD subtypes were driving observed PRS associations with mood-incongruent psychotic features, we did a sensitivity analysis excluding SABD cases. Post-estimation predicted probabilities were plotted to aid interpretation of the PRS associations across RDC subtypes of BD^57^ To correct for multiple comparisons of PRS associations across different phenotypic strata within each model, bootstrapped standard errors and 95% confidence intervals were generated, as an approximation to exact permutation methods ^58^(supplementary E - Note 2). Possible family-wise type-1 error proliferation was controlled for using the Bonferroni Method, calculated by multiplying the bootstrapped p-values by four^59^.

Post-hoc analyses used a MNLM case-control design to examine differential associations across composite phenotypic categories defined by subtype BDI and BD II stratified by psychosis status and a complementary logistic regression analyses comparing the effect of PRS on lifetime occurrence of psychosis, across BD I and BD II subtypes. To examine the distribution of RDC defined cases across levels of PRS, we converted PRS to deciles and generated a stacked bar-chart (SCZ (CLOZUK), SABD, BD I, BD II), by decile.

Analyses were performed using PLINK vl.9 ^60^ or STATA *(Stata Statistical Software: Release 14*. College Station, TX: Stata Corp, LP).

## Results

### Sample description, Genotyping and quality control

After merging BD, CLOZUK and control imputed-genotyped samples and further QC, 18,387 cases and controls (E-supplementary Table 1) with 3,451,354 SNPs with INFO score > 0.8 and MAF >1% were available for analysis. Within the BD sample 52% (N = 2296) of cases endorsed lifetime occurrence of definite psychosis, with <1% missingness in this variable (N=25). Of the BD cases with definite psychosis, 43% (N= 981) were classed as having high lifetime mood-incongruent psychotic features. There was a 9% (N=214) missingness rate for the mood-incongruence variable, within the BD cases with psychosis.

### Case Control PRS associations

As expected (Table 1 Section A), PRS discriminated CLOZUK from controls. PRS in those with a diagnosis of SABD or BD I, but not BD II, were significantly higher than controls.

**Table 1:**
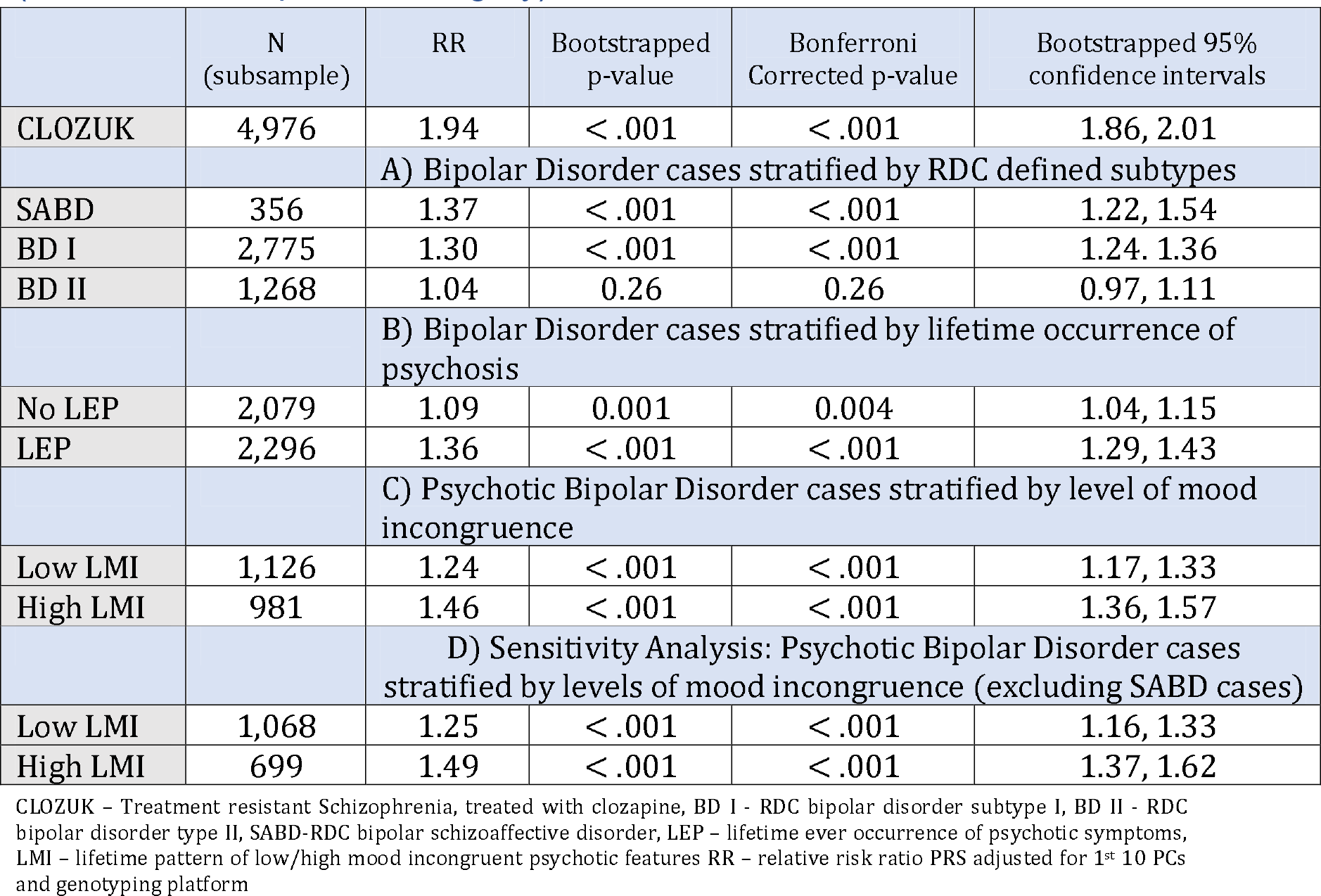
Differential Association of PRS across variously defined BD strata (controls as comparator category)

|  | N<br>(subsample) | RR | Bootstrapped<br>p-value | Bonferroni<br>Corrected p-value | Bootstrapped 95%<br>confidence intervals |
| --- | --- | --- | --- | --- | --- |
| CLOZUK | 4,976 | 1.94 | < .001 | < .001 | 1.86, 2.01 |
| A) Bipolar Disorder cases stratified by RDC defined subtypes |  |  |  |  |  |
| SABD | 356 | 1.37 | < .001 | < .001 | 1.22, 1.54 |
| BD I | 2,775 | 1.30 | < .001 | < .001 | 1.24, 1.36 |
| BD II | 1,268 | 1.04 | 0.26 | 0.26 | 0.97, 1.11 |
| B) Bipolar Disorder cases stratified by lifetime occurrence of psychosis |  |  |  |  |  |
| No LEP | 2,079 | 1.09 | 0.001 | 0.004 | 1.04, 1.15 |
| LEP | 2,296 | 1.36 | < .001 | < .001 | 1.29, 1.43 |
| C) Psychotic Bipolar Disorder cases stratified by level of mood incongruence |  |  |  |  |  |
| Low LMI | 1,126 | 1.24 | < .001 | < .001 | 1.17, 1.33 |
| High LMI | 981 | 1.46 | < .001 | < .001 | 1.36, 1.57 |
| D) Sensitivity Analysis: Psychotic Bipolar Disorder cases stratified by levels of mood incongruence (excluding SABD cases) |  |  |  |  |  |
| Low LMI | 1,068 | 1.25 | < .001 | < .001 | 1.16, 1.33 |
| High LMI | 699 | 1.49 | < .001 | < .001 | 1.37, 1.62 |

CLOZUK – Treatment resistant Schizophrenia, treated with clozapine, BD I - RDC bipolar disorder subtype I, BD II - RDC bipolar disorder type II, SABD-RDC bipolar schizoaffective disorder, LEP – lifetime ever occurrence of psychotic symptoms, LMI – lifetime pattern of low/high mood incongruent psychotic features RR – relative risk ratio PRS adjusted for 1<sup>st</sup> 10 PCs and genotyping platform

### PRS associations within cases

PRS discriminated SCZ from all BD subtypes (Table 2). Within BD, PRS discriminates BD II from both BD I and SABD (Figure 1). The percentage of CLOZUK cases increased monotonically with increasing decile PRS, while the percentage of bipolar subtypes decreased (Figure 2).

**Figure 1.**
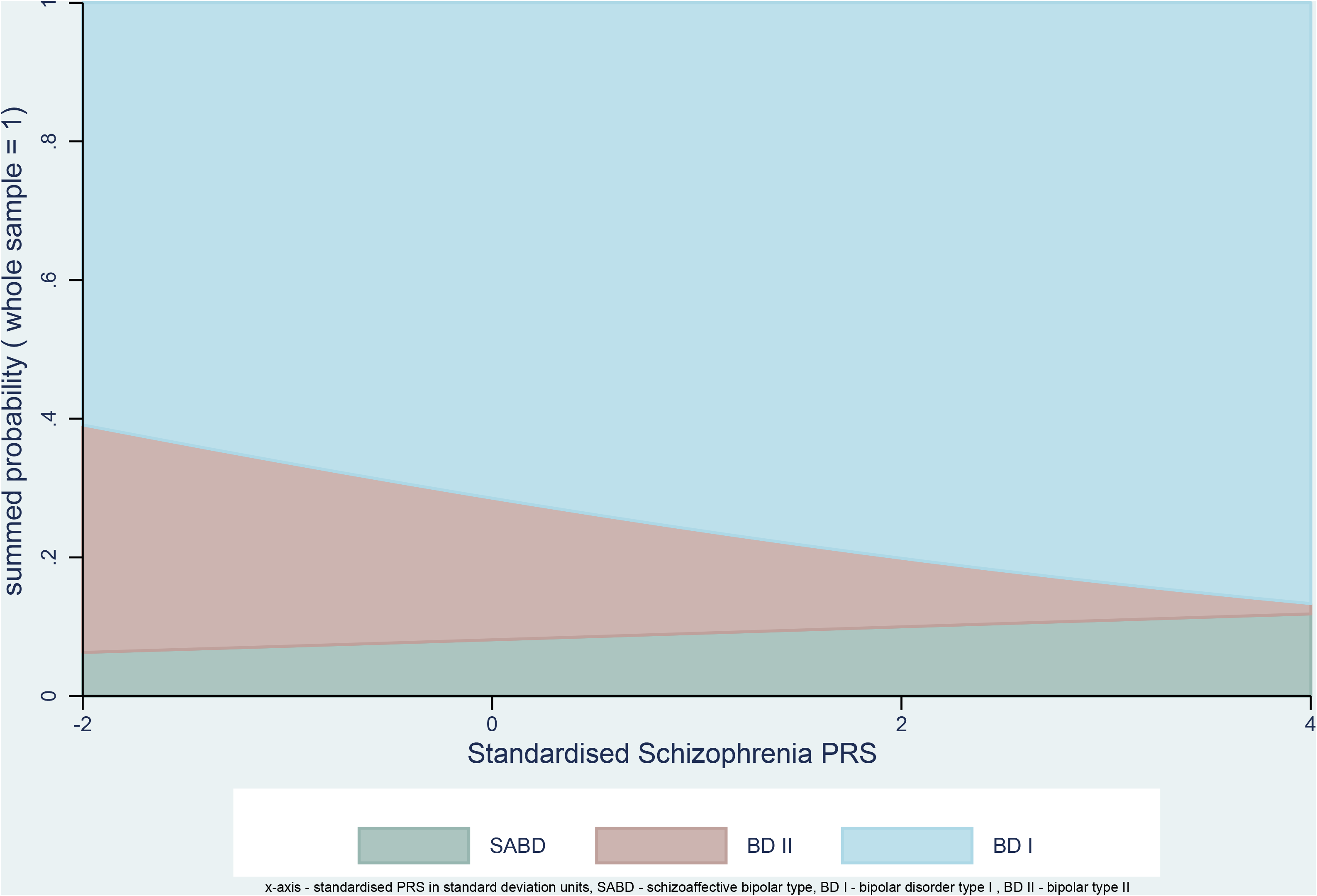
Probability of RDC bipolar subtype as a function of polygenic risk scores (PRS) for schizophrenia

**Figure 2:**
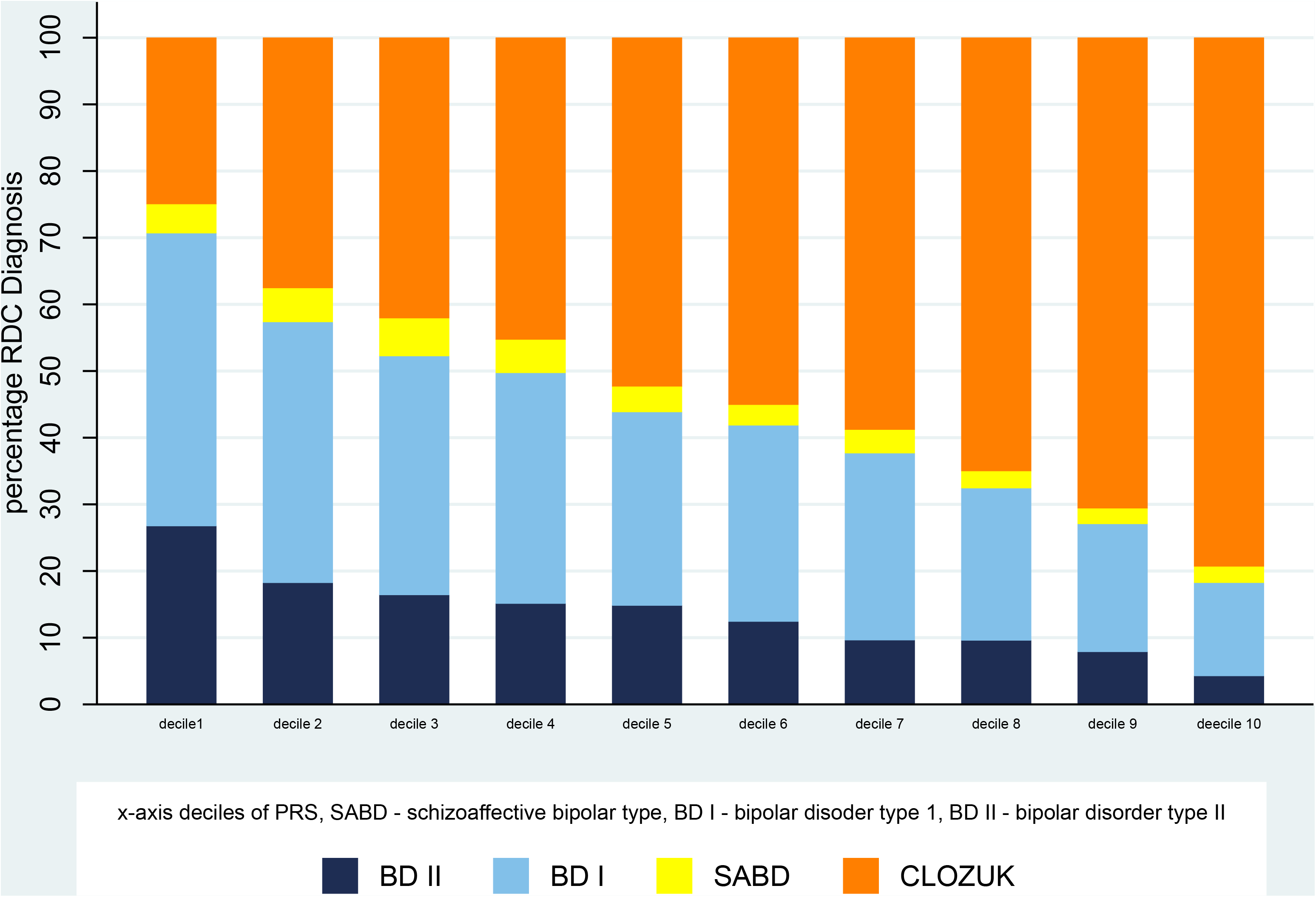
Percentage of bipolar subtype as a function of PRS for schizophrenia - grouped by decile

**Figure 2.**
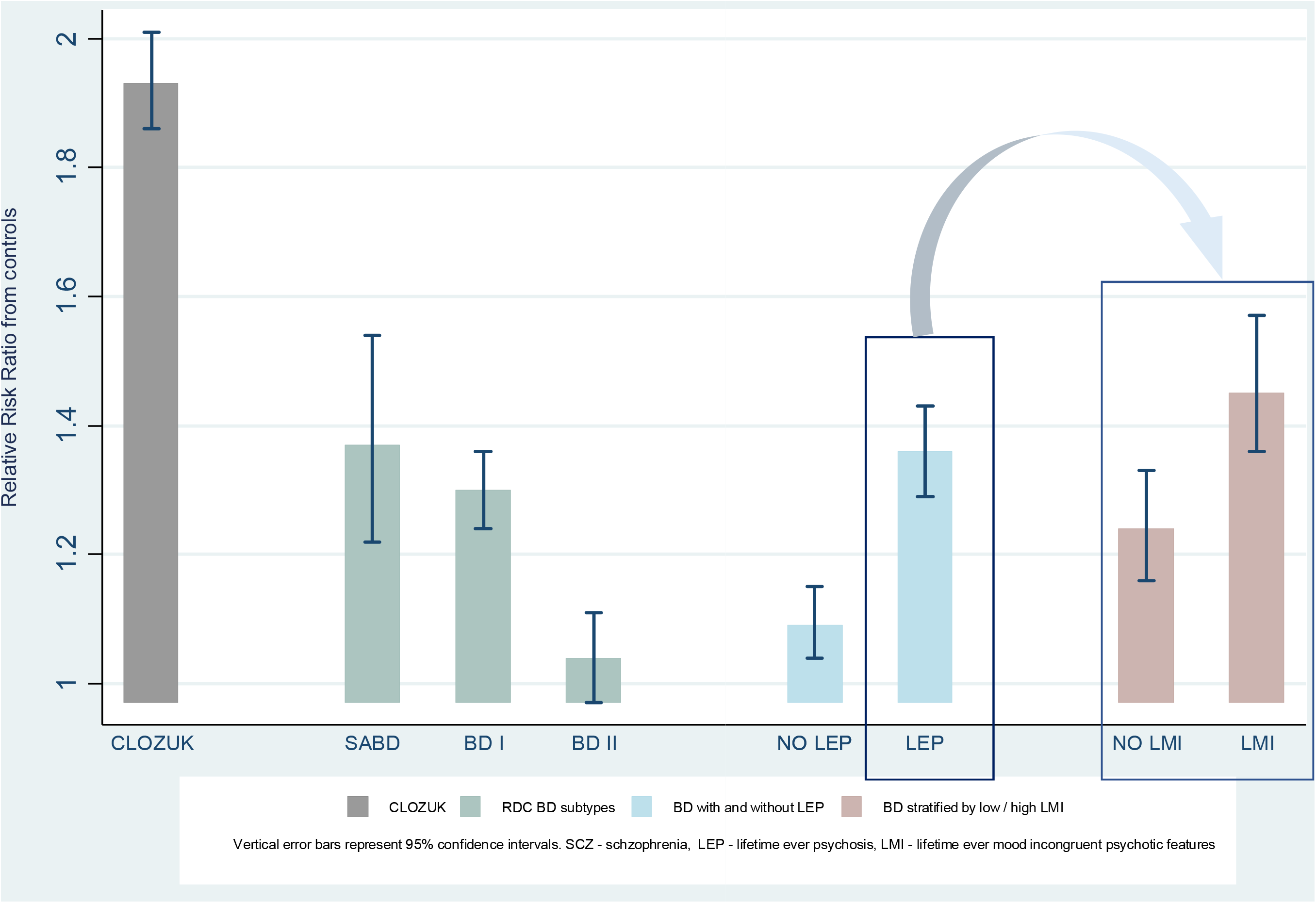
Relative Risk Ratio of PRS with subtypes of BD compared with controls (CLOZUK included for comparison)

**Table 2:** PRS-SCZ associations among cases

|  | RR | Bootstrapped<br>p-value | Bonferroni<br>corrected p-value | Bootstrapped<br>95% C.I. |
| --- | --- | --- | --- | --- |
| SABD compared to TRS | 0.71 | < .001 | < .001 | 0.63, 0.80 |
| BD I compared to TRS | 0.67 | < .001 | < .001 | 0.64, 0.71 |
| BD II compared to TRS | 0.54 | < .001 | < .001 | 0.50, 0.57 |
| SABD compared to BD II | 1.32 | < .001 | < .001 | 1.16, 1.50 |
| BP I compared to BD II | 1.25 | < .001 | < .001 | 1.16, 1.35 |
| SABD compared to BD I | 1.05 | 0.41 | 0.41 | 0.93, 1.18 |

TRS - treatment resistant schizophrenia, treated with clozapine, BDI - RDC bipolar disorder subtype I, BD II - RDC bipolar disorder type II, SABD-RDC bipolar schizoaffective disorder, RR – relative risk ratio PRS adjusted for 1<sup>st</sup> 10 PCs and genotyping platform 95% bootstrapped C.I. - 95% confidence intervals.

### PRS associations with psychotic BD

Compared to controls, the PRS were higher in BD, regardless of whether there was a history of psychosis (Table 1, Section B, Figure 2). However, PRS were significantly higher in BD with psychosis, compared to BD without psychosis (Table 1, Section B, figure 3). Within BD cases, PRS discriminated those with and without psychosis (RR=1.25, 95% bootstrapped adjusted p-value < 001, C.I. (1.16,1.33)).

Post hoc analyses showed the association between PRS and psychosis was present in BD I (OR = 1.21, 95% C.I. 1.10,1.32) but were not statistically significant in BD II (OR = 0.98, 95% C.I. 0.80,1.18). Composite subgroup defined as BD I with psychosis - had higher PRS compared to controls (RR = 1.38,95% C.I. 1.31,1.46) this association was significantly stronger than that of the composite BD I/no psychosis (RR= 1.16, 95% C.I. 1.08,1.25). Within BD II, there was no differential association across subgroups defined by presence/absence of psychosis as compared to controls (supplementary-E: Table-1).

### PRS associations with mood-incongruent psychotic features

Psychotic BD characterised by high mood-incongruence has a higher SCZ polygenic risk burden than controls, with a one standard-deviation increase in PRS increasing the RR of being in the high mood-incongruence category by 46% (RR= 1.46, bootstrapped, 95% C.1. 1.36,1.57) (Figure 3, Table 1 Section C). Although the association was significantly weaker than for the high mood-incongruent group, schizophrenia risk-alleles were enriched in those with low mood-incongruence compared with controls (RR= 1.24, bootstrapped 95% C.I. (1.17, 1.33). Sensitivity analysis excluding the SABD group from analyses found comparable results (Table 1: Section D). Finally, a within-BD-case analysis, measuring mood-incongruence on an ordinal scale found the odds of having higher levels of mood-incongruence, increased with increasing PRS (OR=1.17, (bootstrapped p-value < .001, 95% C.1.1.08 - 1.27)). Analyses excluding the SABD sample found comparable results (0R=1.20, bootstrapped p-value < .001, 95% C.1. 1.09,1.32).

## Discussion

### Main Findings

Higher PRS-SCZ in BD ^20,61^ is well established. Here, we replicate and extend this observation, demonstrating a gradient of PRS associations across SCZ and BD subtypes (CLOZUK > SABD > BD I with psychosis > BD I without psychosis > BD II). We also show BD cases with psychosis carry a higher burden of SCZ risk-alleles, compared to BD without a history of psychosis. Furthermore, individuals with psychotic BD characterised by prominent mood-incongruent psychotic features, carry the highest burden of schizophrenia risk-alleles. There is a clear exposure-response gradient, with increasing PRS associated with psychotic BD and increasing mood-incongruence (mood-incongruent > mood-congruent > no psychosis), supporting our hypothesis that mood-incongruence indexes phenotypic features linked to SCZ liability.

Previously published work examining PRS for SCZ across BD, stratified by psychosis, did not find significant discrimination ^41,62^ although a trend was observed, consistent with the findings presented here. The most likely explanations for the enhanced signal in the current analysis are: PRS were constructed using alleles derived from a larger SCZ-GWAS discovery set which reduces measurement error plus improved power from both this and the larger BD sample ^63^. This group has shown^41^, PRS-SCZ significantly differentiate SABD from non-SABD subtypes, while finding no statistically significant differential between BD stratified by psychosis, suggesting it is the nature of the psychotic symptoms rather than their presence which better indexes liability shared with SCZ. The current analysis supports this proposition that it is the level of mood-incongruence rather than the presence of psychosis *perse* which better specifies a shared biologically-validated dimensional trait, captured, but with less precision by the SABD diagnostic category.

Psychosis and mood-incongruent psychotic features are known to be correlated to poorer prognosis and treatment response^30–32^ It is possible the trans-diagnostic exposure-response gradient for PRS with the occurrence and nature of psychotic symptoms presented here, could be the result of a general psychopathological factor cutting across psychiatric disorders which influences the severity of psychopathology generally, as well as, or rather than a psychosis-specific domain and that PRS derived from SCZ GWAS may be indexing a general liability for psychopathology severity (at least in part)^64^ rather than a (SCZ) disease specific liability.

### Implications

Our study supports the hypothesis that within BD, positive and disorganized psychotic symptoms, and in particular mood-incongruent psychotic features, represent a dimensionally defined stratum with underpinning biological validity. These features are not only phenotypically similar to those observed in prototypal schizophrenia but also index a greater shared genetic aetiology suggesting they share more pathophysiology^65^. It is notable that in those diagnosed with BD I with no history of psychosis, the association with schizophrenia liability was weaker but still on average higher than in the control group, while in the BD II subsample there was no overlap with SCZ liability. We are not suggesting psychotic features are the best or only index of shared pathophysiology, but having established stronger genetic links between the risk for SCZ and BD characterised by the occurrence of psychosis and level of mood-incongruence, we now have a basis to refine this signal. These findings represent a step towards the goal of reconceptualising phenotypic definitions using richer clinical signatures, measured across quantitative/qualitative domains including, symptom loadings and biomarker expression, outlined in the rationale for the Research Domain Criteria (RDoC) ^66,67^ and the road map for mental health research (ROAMER) ^68^ projects. It is probable however a multidimensional stratification process will harness the observed clinical heterogeneity better and define more precise patient-strata/subgroups in closer alignment with the underlying pathophysiology ^68–70^

### Methodological considerations

The phenotypic ratings used in the current analyses are based on both SCAN interviews and case-note review by raters with excellent inter-rater reliability, which is expected to minimise rates of missing data and reduce the likelihood of phenotypic misclassification^71^. Our psychosis phenotypes are broadly defined and likely to represent imperfect measurements of a continuously distributed phenotype^72^, imposing categorical constraints as we have done may reduce power. We generated PRS using a single discovery set p-value threshold < 0.05 and dealt with multiple comparisons, across different phenotypic categories/strata using bootstrap re-sampling approaches within each of our 4 independent analyses, adjusting for family-wise type-1 error proliferation using Bonferroni’s correction. We have mitigated against potential confounding due to population stratification and potential batch effects across cases and controls, by partialling out the first 10 PCs and genotyping platforms from the PRS. The PRS were generated using most probable genotypes which can potentially reduce power due to a small (non-differential) loss of information at some markers making our results conservative, but the conclusions are unlikely to change. Finally, we have only examined the effect of common variants, as rare variants are not captured by current GWAS.

## Conclusions

We show for the first time a gradient of polygenic liability across schizophrenia and bipolar disorder, indexed by the occurrence and level of mood-incongruence of positive and disorganised psychotic symptoms. This highlights the usefulness of genetic data to dissect clinical heterogeneity within and across disorders, and suggests further research could potentially aid in defining patient stratifiers with improved biological precision/validity, moving us tentatively towards precision medicine in psychiatry.

## Acknowledgements

The work at Cardiff University was funded by Medical Research Council (MRC) Centre (MR/L010305/1) and Program Grants (G0800509). The CLOZUK sample was genotyped with funding from the European Union’s Seventh Framework Programme for research, technological development and demonstration under grant agreement n° 279227 (CRESTAR Consortium; http://www.crestar-proiect.euA). For the CLOZUK sample we thank Leyden Delta (Nijmegen, Netherlands) for supporting the sample collection, anonymisation and data preparation (particularly Marinka Helthuis, John Jansen, Karel Jollie and Anouschka Colson) and Andy Walker from Magna Laboratories (UK). We acknowledge Lesley Bates, Catherine Bresner and Lucinda Hopkins, at Cardiff University, for laboratory sample management

We would like to acknowledge funding for BDRN from the Wellcome Trust and Stanley Medical Research Institute, all members of BDRN, and especially the participants who have kindly given their time to participate in our research.

Generation Scotland (GS) received core support from the Chief Scientist Office of the Scottish Government Health Directorates [CZD/16/6] and the Scottish Funding Council [HR03006]. Genotyping of the GS:SFHS samples was carried out by the Genetics Core Laboratory at the Wellcome Trust Clinical Research Facility, Edinburgh, Scotland and was funded by the Medical Research Council UK and the Wellcome Trust (Wellcome Trust Strategic Award "STratifying Resilience and Depression Longitudinally” (STRADL) Reference 10403 6/Z/14/Z)

The Type 1 Diabetes Genetics Consortium (T1DGC; EGA dataset EGAS00000000038) is a collaborative clinical study sponsored by the National Institute of Diabetes and Digestive and Kidney Diseases (NIDDK), National Institute of Allergy and Infectious Diseases (NIAID), National Human Genome Research Institute (NHGRI), National Institute of Child Health and Human Development (NICHD), and JDRF. Venous blood collection for the 1958 Birth Cohort (NCDS) was funded by the UK’s Medical Research Council (MRC) grant G0000934, peripheral blood lymphocyte preparation by Juvenile Diabetes Research Foundation (JDRF) and WT and the cell-line production, DNA extraction and processing by WT grant 06854/Z/02/Z. Genotyping was supported by WT (083270) and the European Union (EU; ENGAGE: HEALTH-F4-2007-201413).

The funders have had no role in the design and conduct of the study; collection, management, analysis, and interpretation of the data; preparation, review, or approval of the manuscript; and decision to submit the manuscript for publication

V Escott-Price had full access to all the data in the study and takes responsibility for the integrity of the data and the accuracy of the data analysis.

J. Allardyce, G. Leonenko, A F J. Pardiñas, M. Hamshere and V. Escott-Price, all from MRC Centre for Neuropsychiatric Genetics and Genomics, Institute of Psychological Medicine and Clinical Neurosciences, School of Medicine, Cardiff University, Cardiff, UK conducted and are responsible for the data analysis.

